# DAF-16/Foxo suppresses the transgenerational sterility of *prg-1* piRNA mutants via a systemic small RNA pathway

**DOI:** 10.1101/326751

**Authors:** Matt Simon, Maya Spichal, Bree Heestand, Stephen Frenk, Ashley Hedges, Malik Godwin, Alicia S. Wellman, Aisa Sakaguchi, Shawn Ahmed

**Affiliations:** Department of Genetics, University of North Carolina, Chapel Hill, NC, USA; Department of Biology, University of North Carolina, Chapel Hill, NC, USA; Curriculum in Genetics and Molecular Biology, University of North Carolina, Chapel Hill, NC, USA; University of Rochester, NY, USA; University of Utah Hospital, Salt Lake City, UT, USA; Graduate School of Science, Osaka University, Osaka, Japan

**Author notes:** These authors contributed equally to this work (Matt Simon, Maya Spichal, Bree Heestand, Stephen Frenk).

## Abstract

Mutation of the *daf-2* insulin/IGF-1 receptor activates the DAF-16/Foxo transcription factor to suppress the transgenerational sterility phenotype of *prg-1*/piRNA mutants that are deficient for piRNA-mediated genome silencing. As with PRG-1/piRNAs, mutations in the nuclear RNA interference gene *nrde-1* compromised germ cell immortality, but deficiency for *daf-2* did not suppress the transgenerational sterility of *nrde-1* or *nrde-4* single mutants or of *prg-1; nrde-4* or *prg-1; hrde-1* double mutants. NRDE-1 and NRDE-4 promote transcriptional silencing in somatic cells via the nuclear Argonaute protein NRDE-3, which was dispensable for germ cell immortality. However, *daf-2* deficiency failed to promote germ cell immortality in *prg-1; nrde-3* mutants. Consistently, we found that DAF-16 activity in somatic cells suppressed the transgenerational sterility of *prg-1* mutants via the SID-1 dsRNA transmembrane channel that promotes systemic RNAi as well as Dicer, the dsRNA binding protein RDE-4 and the RDRP RRF-3. We conclude that DAF-16 activates a cell-non-autonomous systemic RNAi pathway that promotes small RNA-mediated genome silencing in germ cells to suppress loss of the genomic immune surveillance factor Piwi/PRG-1.

**Author Summary:** Small RNAs can promote genome silencing. The Argonaute protein Piwi interacts with thousands of small RNAs termed piRNAs in germ cells to suppress expression of transposons and foreign genetic elements. However, the Piwi silencing system may be commonly targeted by viral or transposon genomic parasites that seek to suppress the endogenous defences against their expression and replication. Activation of the DAF-16 stress response pathway promotes adult longevity and can also abolish the transgenerational sterility of *C. elegans* Piwi mutants. We found that DAF-16 accomplishes this by activating a somatic small RNA pathway where small RNAs are initially produced in the soma and are then transported into the germline to suppress expression of a toxic genetic locus in Piwi mutants. Thus, the DAF-16 stress response pathway activates a systemic small RNA cascade to suppress defects in the Piwi/piRNA genome silencing system.

## Introduction

Small RNAs are key regulators of the transcriptome in eukaryotes, regulating RNA stability or translation in the cytoplasm and transcription in the nucleus (1). Small RNA-mediated transcriptional silencing in germ cells of metazoans is carried out by the Piwi Argonaute protein and Piwi-associated small RNAs termed piRNAs, which suppress expression of transposable elements and other genomic parasites (2). This has several consequences. First, Piwi promotes genome stability by suppressing the potentially mutagenic effects of transposition or viral integration, which could have toxic, neutral or beneficial effects (3–5). Second, it promotes heterochromatin formation at pericentromeres, which are transposon-rich segments of the genome that create large blocks of constitutive heterochromatin in order to promote accurate chromosome segregation during mitosis (6). Aside from its silencing properties, *Drosophila* Piwi can also promote transcriptional activation, as can the *C. elegans* anti-silencing Argonaute protein CSR-1 (7, 8). *Drosophila* Piwi can also regulate HSP90 activity and can promote mitotic stem cell function and meiosis (7, 9).

The *C. elegans* Piwi ortholog PRG-1 interacts with piRNAs to promote silencing of transposons and some genes. *prg-1* mutants display activation of the Tc3 DNA transposon (10, 11), but the overall frequency of de novo transposition events in *prg-1* mutants is very low (12). This is because *C. elegans* PRG-1/Piwi can act to initiate silencing of transposons or transgenes that enter the germ line, but genomic silencing is then largely maintained by Mutator proteins that utilize RNA dependent RNA polymerases (RDRPs) to promote the biogenesis of endogenous siRNAs which are 22 nucleotides long and have a G as their first nucleotide, referred to as 22G-RNAs (13, 14). Loss of Mutator-mediated 22G RNA biogenesis leads to significant levels of de novo transposition, termed a Mutator phenotype (15–17). Therefore, although PRG-1/piRNAs act in perinuclear P-granules in conjunction with neighbouring Mutator bodies to promote the biogenesis of 22G RNAs that promote genomic silencing in response to foreign nucleic acids (18), this silencing is often maintained in a manner that is independent of PRG-1/piRNAs or only partially dependent on PRG-1 (19). 22G-RNAs that are created in response to piRNAs can associate with the germline-specific nuclear Argonaute protein HRDE-1 to promote silencing of foreign transgenes (13, 14, 20). HRDE-1 and associated 22G RNAs promote genome silencing in germ cells via the Nuclear RNAi Defective (NRDE) proteins NRDE-1, NRDE-2 and NRDE-4 that stimulate the activities of histone modifying enzymes to cause genomic silencing in response to small RNAs (20–22). The nuclear RNAi pathway can target foreign transgenes for genomic silencing. Once established, this silent state can persist for many generations in the absence of piRNAs, termed RNAi inheritance (13, 14, 20, 23, 24).

Piwi mutants in many species are immediately sterile (7), whereas *C. elegans prg-1* mutants are fertile but have been observed to display reduced levels of fertility at elevated temperatures (10, 11, 25). Outcrossed *prg-1* mutants initially display robust levels of progeny but after many generations will become completely sterile, indicating that PRG-1/Piwi promotes germ cell immortality (12). It is unlikely that the very modest levels of transposition in *prg-1* mutants (10, 11) are responsible their defect in germ cell immortality, as *mutator* mutants that display significant levels of transposon activity in germ cells remain fertile indefinitely if grown at low temperatures (12).

dsRNA can be administered to *C. elegans* either through uptake from the gut by E. coli that they feed on, direct injection, soaking, or expression of hairpin-forming transgenes or through uptake from the gut by *E. coli* that they feed on (26). This exogenous RNAi pathway requires the Dicer nuclease DCR-1 and its dsRNA binding cofactor RDE-4 (1). RNAi-induced in response to exogenous dsRNA triggers can spread throughout the animal and be passed to the next generation via germ cells (27). Similar to the transgene silencing above, RNAi-induced silencing of endogenous loci can persist for many generations (28). Transmission of RNAi between cells depends on the Systemic Interference Defective (Sid) genes, which include the SID-1 transmembrane protein that promotes import of short dsRNA fragments into cells (29, 30). The systemic RNAi pathway can respond to dsRNAs expressed in somatic cells to induce a persistent heterochromatic genome silencing over many generations (31). Endogenous functions of this systemic RNAi pathway are not well understood but are likely to include viral immunity (32, 33).

A second Dicer complex, the Enhanced RNAi (ERI) complex, is required for silencing of endogenous loci, including duplicated genes and transposable elements, which are silenced independently of the piRNA pathway (34–36). The ERI complex generates a cohort of primary small RNAs termed 26G RNAs, utilising the RNA dependent RNA polymerase RRF-3 (37, 38). Both classes of primary small RNAs generated by Dicer are amplified via RDRPs into effector 22G-RNA populations that bear perfect homology to their targets to promote either transcriptional or post-transcriptional silencing (16). In the absence of the 26G RNA Eri pathway, the response to exogenous RNAi is enhanced (39), likely due to release of limiting amounts of proteins such as DCR-1 and RDE-4, which are components of both exogenous RNAi and endogenous 26G pathways (37, 38).

Although deficiency for *prg-1*/piRNAs leads to complete sterility after growth for many generations, bouts of starvation can delay the onset of sterility (12). Intriguingly, mutation of the *daf-2* insulin/IGF1 receptor (40), whose activity is suppressed by starvation, strongly suppresses the transgenerational fertility defect of p*rg-1* mutants (12). The DAF-16 transcription factor that functions downstream of *daf-2* to promote longevity and stress resistance is required for the ability of starvation and *daf-2* mutation to ameliorate the fertility defects of *prg-1* mutants (12). The exact cause of the transgenerational sterility of *prg-1* mutants and how this is suppressed by *daf-2* mutations are unknown. However, the suppression of sterility by *daf-2* mutation requires the activity of the endogenous RNAi pathway as it requires 22G-RNAs and H3K4 demethylases that promote genomic silencing (12). These results suggest that *daf-2* acts to compensate for *prg-1* deficiency by upregulating an alternative small RNA pathway that silences repetitive regions of the genome.

Here we demonstrate that DAF-16 suppresses the fertility defects of *prg-1* by activating a systemic small RNA pathway that requires somatic and germline nuclear Argonaute proteins, as well as nuclear RNAi factors that function downstream of PRG-1/piRNAs to promote germline immortality. Therefore, an endogenous function of systemic RNAi in *C. elegans* is to promote genomic silencing in the germline if the Piwi/piRNA germline immune response system is compromised.

## Results

### A nuclear silencing process promotes germ cell immortality

*tir-1* encodes a toll-like receptor and is required for innate immunity (41–43). Strains containing the *tir-1(ky388)* mutation displayed sterility after growth for a few generations but can be rejuvenated after outcrosses with N2 wildtype (C. Bargmann, personal communication). This suggested a defect in germ cell immortality, but strains deficient for independent mutations in *tir-1*, *tm1111* and *tm3036*, failed to become sterile (n=2 strains per allele, 30 generations of growth), suggesting that *tir-1(ky388)* contained a linked mutation responsible for its Mortal Germline phenotype. Three-factor crosses with *+ tir-1(ky388) + / unc-93 + dpy-17* heterozygotes revealed a novel *mortal germline* mutation *yp4* located just to the left of *tir-1* (Fig 1A,C-D, S1 Table).

**Fig 1.**
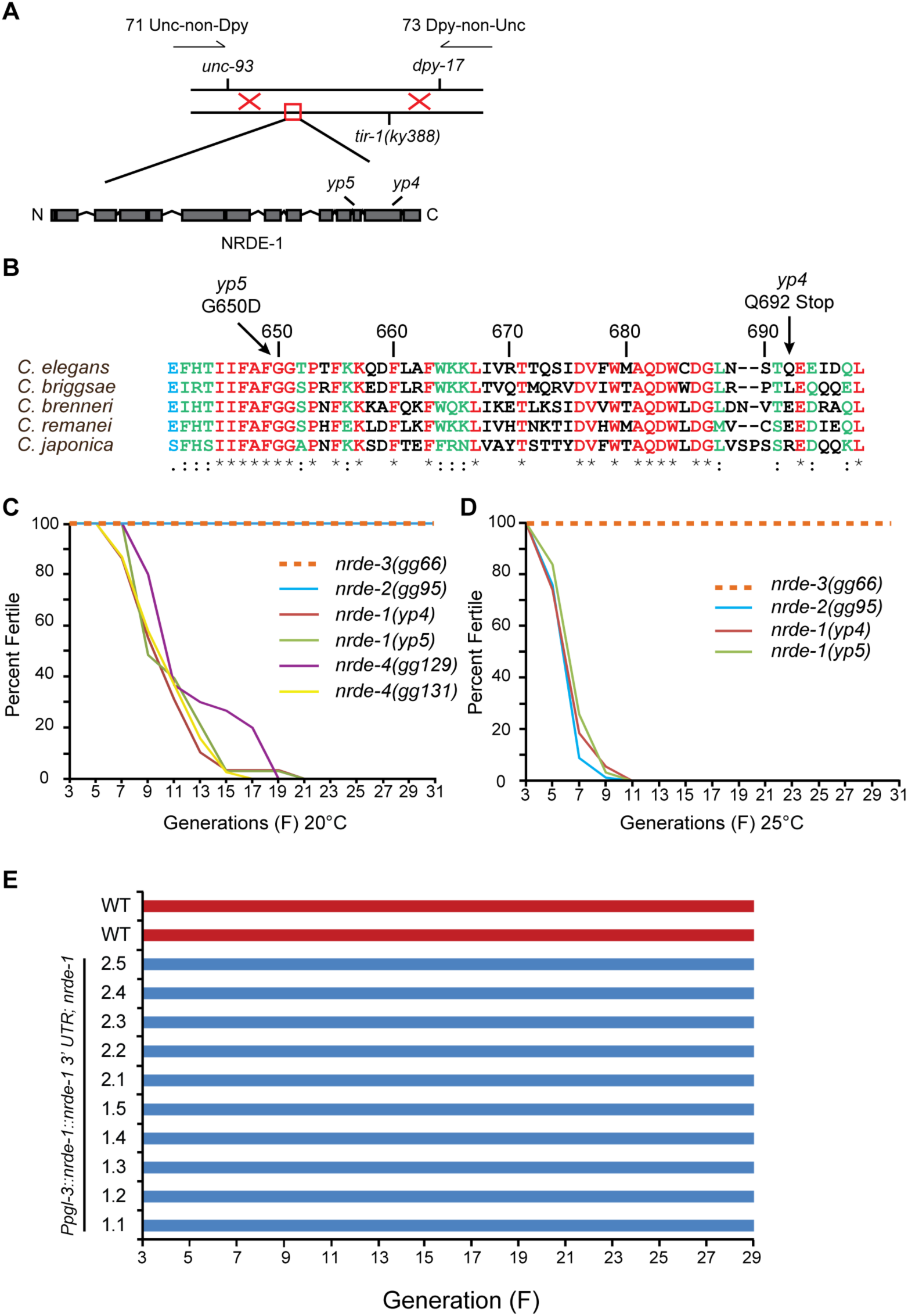
*nrde-1, nrde-2* and *nrde-4* promote germ cell immortality. (A) Three-factor mapping of *nrde-1* revealed a genetic map position at -4.6 on Chromosome *III*. Sequence analysis of this region of the genome revealed mutations in the gene C14B1.6 for two *nrde-1* alleles. (B) Alignment of peptides for NRDE-1 homologs of five nematode species. *yp4* and *yp5* mutations annotated with arrows. (C) Progressive sterility of *nrde-1* and *nrde-4* at 20°C. *nrde-2* and *nrde-3* showed no transgenerational sterility. (D) *nrde-1* and *nrde-2*, but not *nrde-3*, showed progressive sterility at 25°C.

Analysis of ethylmethane sulphonate-induced *mrt* mutants revealed one mutation, *yp5*, with a map position similar to that of *yp4*. *unc-93 yp4 + / + yp5 dpy-17* but not *unc-93 + + / + yp5 dpy-17* or *unc-93 yp4 + / + + dpy-17* trans-heterozygotes resulted in progressive sterility (n=4, 5 and 6 strains propagated, respectively) (Fig 1A, S1 Table), indicating that *yp5* fails to complement *yp4* and that their Mrt phenotypes are likely to be caused by mutations in the same gene. Sequencing of ∼120 kb in the genetic map interval of *yp4* revealed 4 mutations in *yp4* mutant genomic DNA, one of which caused a stop codon at amino acid Q692 in the predicted protein product of C14B1.6 (Fig 1B). Genomic DNA of *yp5* possessed a G to T point mutation that in C14B1.6 is predicted to cause a G650D substitution in an amino acid that is conserved in C14B1.6 homologs in closely related nematodes (Fig 1B). BLAST analysis failed to reveal homologs of this protein in more distantly related organisms. While this work was in progress, C14B1.6 was defined to encode the nuclear RNA interference gene *nrde-1*, a nuclear RNAi protein that promotes histone methylation at loci targeted for silencing by exogenous dsRNA triggers (21).

Most nuclear RNAi proteins are likely to be ubiquitously expressed in *C. elegans* tissues and function in both germ and somatic cells (20–22). Independent single copy insertions of *Ppgl-3::nrde-1::nrde-1 3’UTR* (44), a *nrde-1* transgene driven by the germline-specific promoter of *pgl-3* (45), rescued the Mortal Germline phenotype of *nrde-1(yp4)*, indicating that NRDE-1 functions in germ cells to promote germ cell immortality (Fig 1C-E).

To further understand the role of nuclear RNAi in germline immortality, we outcrossed mutations in the remaining components of the Nrde pathway, which promote transcriptional silencing in somatic cells in response to exogenous dsRNAs, and tested these for defects in germ cell immortality. Like NRDE-1, deficiency for a second nuclear RNAi protein with no homologs outside of closely related nematodes, NRDE-4 (21), resulted in progressive sterility at 20°C (average 50% transgenerational lifespan of 10.22 generations n=4 experiments) (Fig 1C, 2A-B, S1 Table). *nrde-2* mutants remained fertile indefinitely at 20°C but became progressively sterile if propagated at 25°C (Fig 1C-D, S1 Table) (22), whereas deficiency for the somatic Argonaute protein required for nuclear RNAi, NRDE-3 (46), resulted in strains that could be propagated indefinitely at any temperature (Fig 1C-D, S1 Table). However, deficiency for the germline nuclear Argonaute protein HRDE-1 (20) led to progressive sterility only at 25°C (Fig 2C, S1 Table). In conclusion, the germline components of the nuclear RNAi pathway, NRDE-1, NRDE-2, NRDE-4 and HRDE-1, promote germ cell immortality. While this study was in progress, analogous conditional and non-conditional defects in germ cell immortality were reported for strains with defects in the nuclear silencing proteins HRDE-1, NRDE-1, NRDE-2 and NRDE-4 mutants (20).

**Fig 2.**
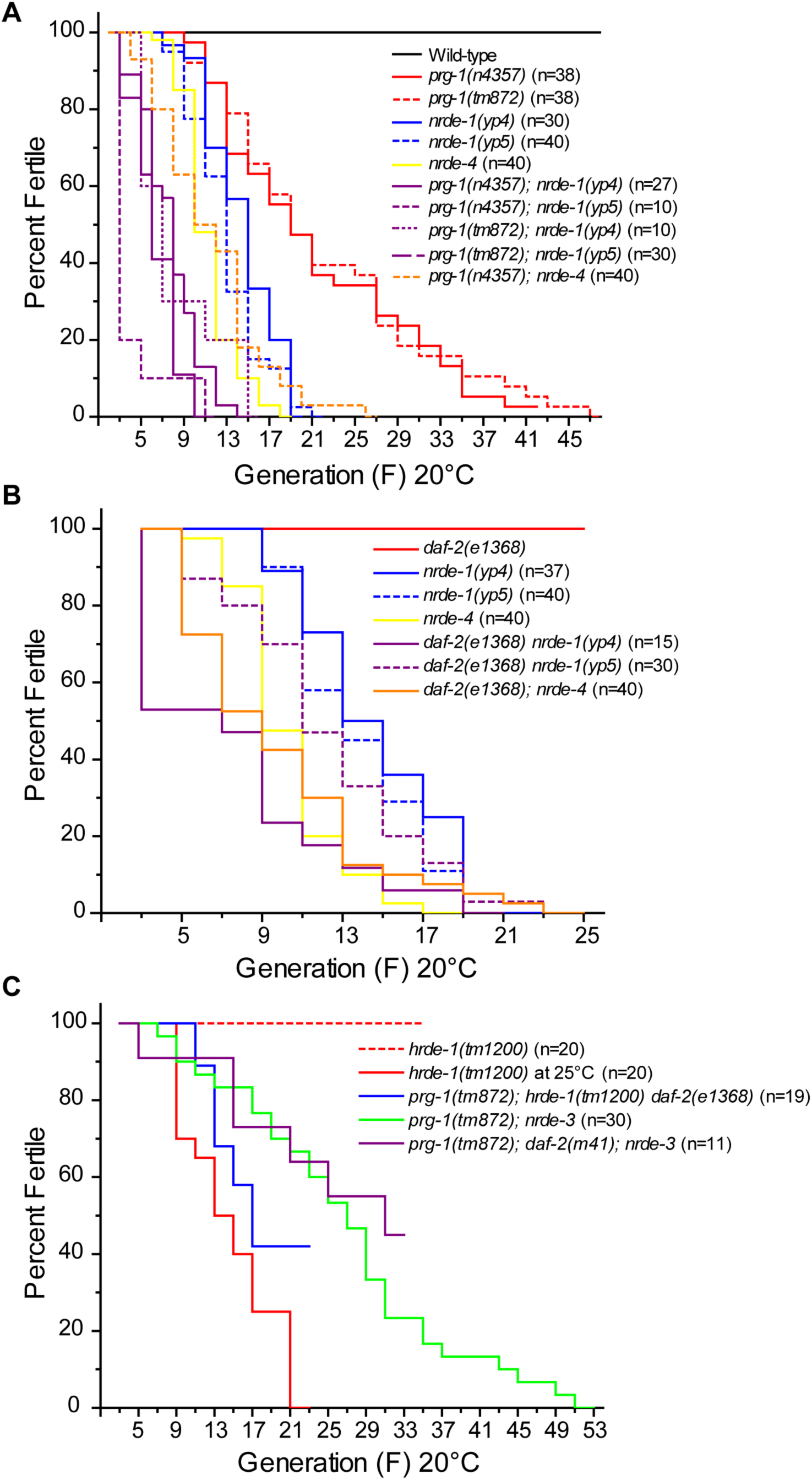
*nrde* factors are required for *prg-1; daf-2* germ cell immortality. Mortal Germline assays for (A) *prg-1; nrde* and control strains, (B) *daf-2; nrde* and control strains, and (C) *prg-1; hrde-1 daf-2* (blue)*, prg-1; daf-2; nrde-3* (purple) and control strains. Survival analysis in S1 Table. P values are provided in S2 Table.

### *nrde* factors are required for *daf-2* to suppress the fertility defect of *prg-1*

The germline nuclear Argonaute protein HRDE-1, and the nuclear RNAi factors NRDE-1, NRDE-2 and NRDE-4 act downstream of PRG-1/piRNAs to promote dsRNA-induced inheritance of transgene silencing in germ cells over many generations. Therefore, we asked if the Mortal Germline phenotypes of *prg-1* and *nrde-1* or *nrde-4* are additive. *prg-1* single mutants had a significantly longer transgenerational lifespan when compared to either *nrde-1 or nrde-4* (p<0.003 all comparisons, Fig 2A, S2 Table). When *prg-1* was combined with *nrde-4*, the transgenerational sterility phenotypes of these double mutants were not significantly shorter than *nrde-4* single mutant controls (p=1, Fig 2A, S2 Table). Additionally, when *prg-1* was combined with *nrde-1*, the transgenerational lifespan was not like *prg-1*, but short like *nrde-1*, often with *prg-1; nrde-1* double mutants having shorter transgenerational lifespans than *nrde-1* or *nrde-4* single mutants (Fig 2A, S1 Table, S2 Table). We conclude that although *prg-1* and the *nrde* genes promote epigenetic silencing in germ cells and germ cell immortality, that *nrde* genes may function at least partially in parallel with *prg-1* in maintaining germ cell immortality. Consistently, PRG-1/Piwi and the NRDE genes have recently been suggested to silence partially overlapping segments of the genome, for example transposon classes, possibly due to a form of redundancy that has been built into the *C. elegans* genome silencing system (47).

Given that *prg-1* deficiency can be suppressed by *daf-2*, we combined *daf-2* with *nrde-1* or *nrde-4*, to determine if *daf-2* can suppress the fertility defect of the *nrde* mutants. However, *daf-2; nrde* double mutants still became sterile similar to *nrde* single mutant controls, as did *prg-1; daf-2; nrde-4* triple mutants (Fig 2B, Fig S1, S1 Table, S2 Table). We also tested the germline nuclear Argonaute protein that functions upstream of NRDE-1 and -4 to promote transgene silencing in the germline, HRDE-1 (13, 14, 20, 24). Given that *hrde-1* mutants do not become sterile at 20°C (Fig 2C) and given that *daf-2* mutants arrest as dauer larvae at 25°C (48), we created *prg-1; daf-2 hrde-1* triple mutants and found that these became sterile (Fig 2C, S1 Table). Therefore, the germline silencing function of the nuclear RNAi pathway is required for *daf-2* to suppress the fertility defects of *prg-1* mutants. We next tested the somatic branch of the *nrde* pathway using NRDE-3, a nuclear Argonaute protein that is specific to somatic cells and is not required for germ cell immortality (Fig 1C-D, S1 Table). We found that *prg-1; daf-2; nrde-3* triple mutants displayed transgenerational sterility similar to that of *prg-1; nrde-3* (p=1, Fig 2C, S1 Table, S2 Table). Therefore, both somatic and germline branches of the nuclear RNAi pathway are required for *daf-2* deficiency to suppress the fertility defects of *prg-1*. We further analysed this somatic RNAi pathway by characterizing GFP::NRDE-3 expression (46) in embryos of wild-type, *prg-1*, *daf-2* and *prg-1; daf-2* mutants. While all wild-type and *prg-1* mutants displayed GFP::NRDE-3 mainly in the nucleus, *daf-2* and *prg-1; daf-2* mutants showed a more frequent localization of GFP::NRDE-3 to the cytoplasm and an absence of clear nuclear localization for the same embryonic stages (Fig 3A-B). Given that NRDE-3::GFP becomes more nuclear in response to exogenous dsRNA triggers that induce nuclear silencing (46), this suggests that a compromised DAF-2/insulin/IGF1 pathway may influence endogenous levels of nuclear small RNA silencing in the soma.

**Fig 3.**
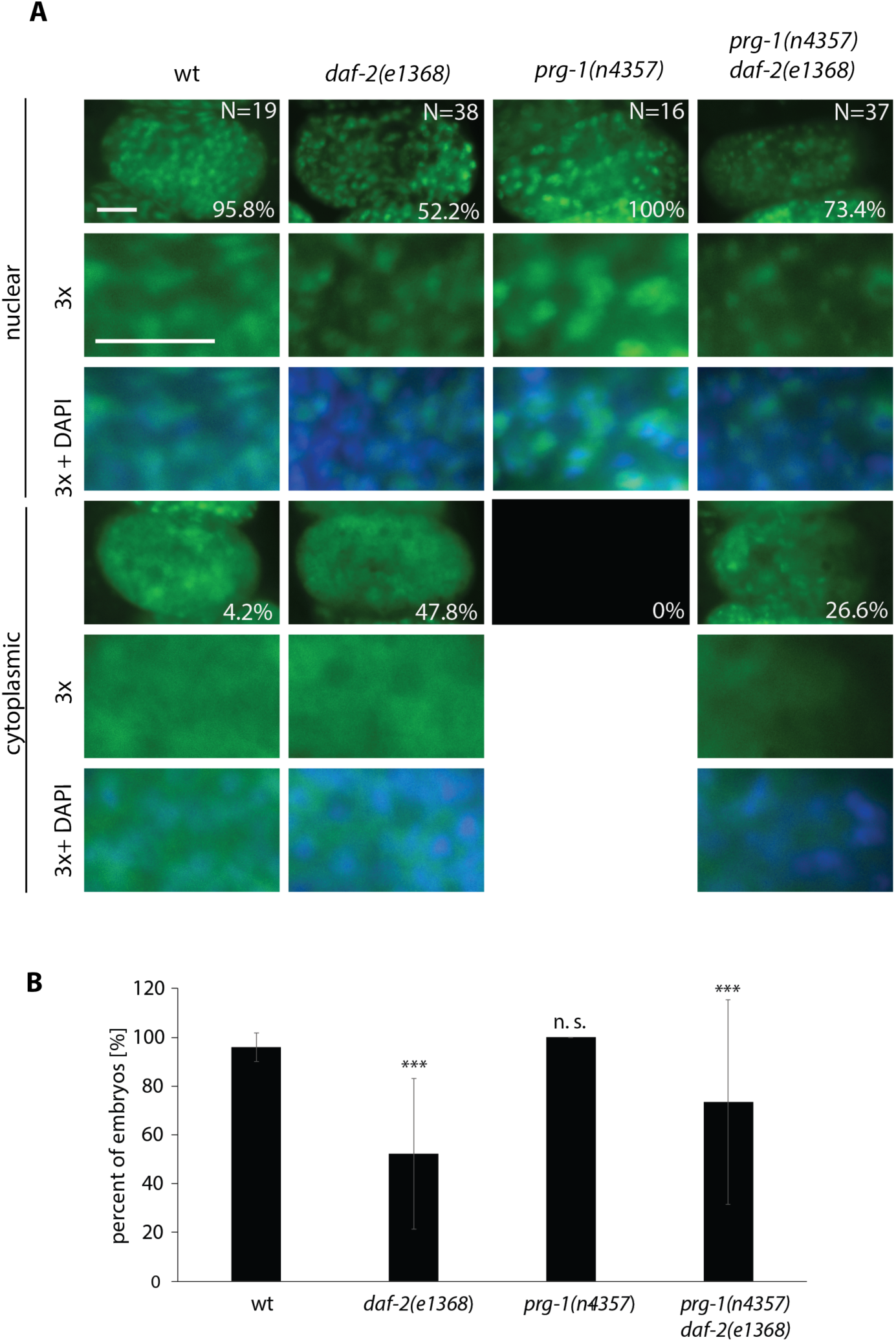
Embryonic expression of GFP::NRDE-3. (A) Images of *C. elegans* embryos at 110 -190 cell stage expressing GFP::NRDE-3 with indicated genotypes. Nuclear and mostly cytoplasmic GFP::NRDE-3 expression is depicted with a 3-fold magnification and percentages of embryos expressing mostly either phenotype is indicated. (B) Quantification of embryos expressing clear nuclear GFP::NRDE-3. Error bars are expressed as standard deviation and significance was determined using a chi-square test.

### DAF-16 acts in somatic cells to suppress *prg-1*

When *daf-2* is deficient, the DAF-16 transcription factor functions in a cell-non-autonomous manner in somatic cells to promote longevity (49, 50). To better characterize the mechanism whereby the DAF-16 pathway suppresses the fertility defect of *prg-1* mutants, we tested independent repetitive DAF-16 transgenes that rescue the longevity defect caused by deficiency for *daf-16*, *muIs71 [GFP::daf-16a (bKO)]*, which was constructed from a large genomic clone of the *daf-16* locus (50), and a second transgene *zIs356 [daf-16a::GFP]* that contains the cDNA for *daf-16a* fused to *GFP* (51). We found that 4 combinations of *prg-1 daf-16; daf-2* strains containing either *daf-16 transgene* could be propagated indefinitely, as observed for *prg-1; daf-2* mutants (6 allele combinations, p=1 all comparisons, Table S2) and in stark contrast to the progressive sterility phenotype of *prg-1 daf-16; daf-2* strains (8 allele combinations, p<0.0002 all comparisons, Table S2) (Fig 4A, S1 Table, S2 Table).

**Fig 4.**
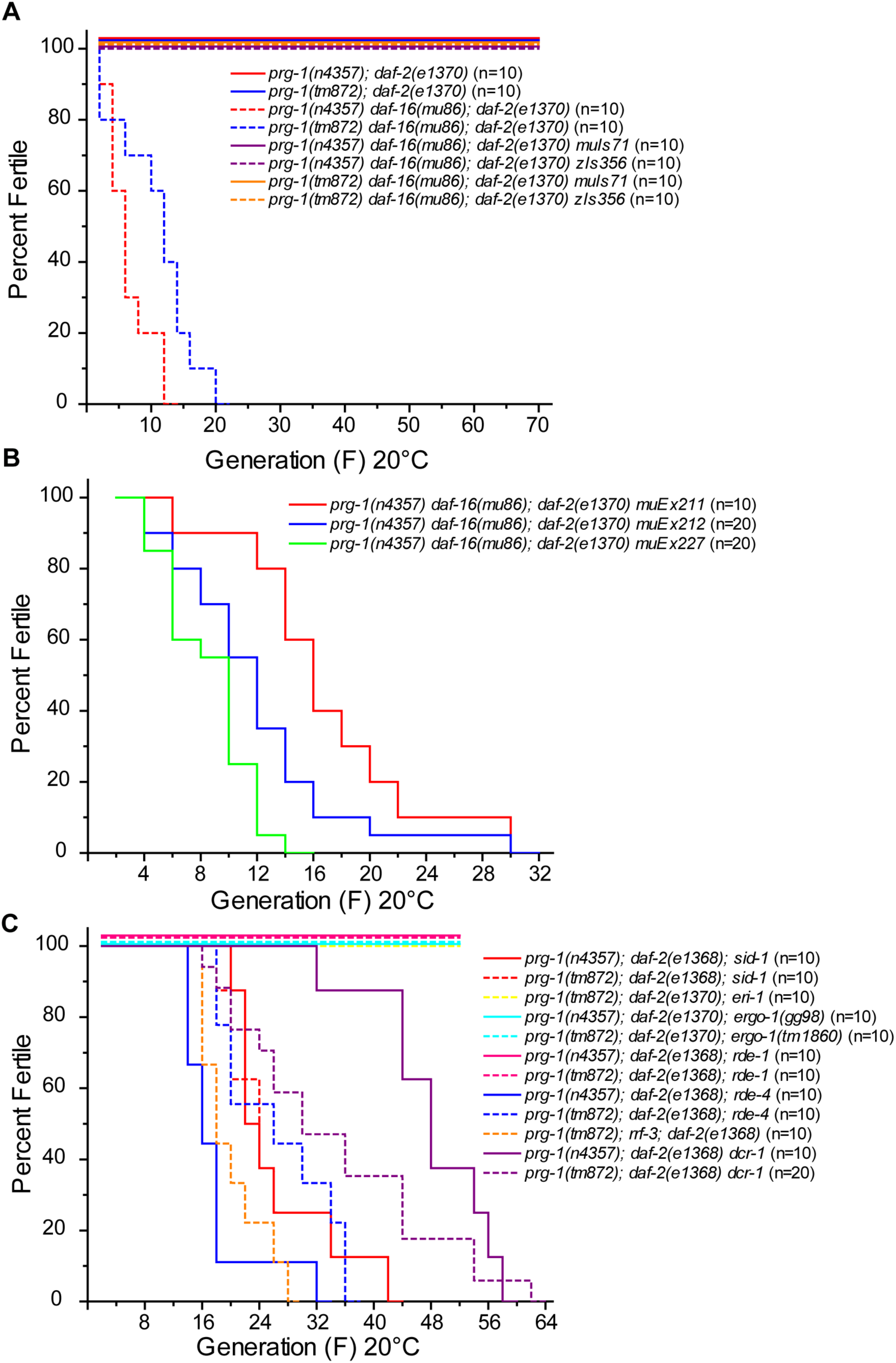
Somatic DAF-16 suppresses prg-1 germline immortality. Mortal Germline assays for (A) *prg-1 daf-16; daf-2 muIz71* or *zIs356* and controls, (B) *prg-1 daf-16; daf-2; muEx211* or *prg-1 daf-16; daf-2; muEx212* or *prg-1 daf-16; daf-2; muEx227*, and (C) *prg-1; daf-2* combined with several RNAi pathway mutations. Survival analysis in S1 Table. P values are provided in S2 Table.

*muIs71* and *zIs356* are repetitive transgenes that are commonly silenced in germ cells via ‘cosuppression’ (52, 53), a small RNA-based phenomenon that silences both repetitive transgenes and as well as endogenous loci that are homologous to the transgene and foreshadowed the discovery of RNA interference (54, 55). PRG-1/piRNAs play roles in initiation and in some cases maintenance of transgene silencing (13, 14, 19, 24). We therefore sought to confirm the model that somatic DAF-16 is responsible for the mechanism by which *daf-2* mutation suppresses the transgenerational fertility defect of *prg-1*. We found that animals that solely contain the *zIs356* DAF-16 transgene fail to express DAF-16::GFP in germ cells of otherwise wildtype animals or of *prg-1 daf-16* and *prg-1 daf-16, daf-2* mutant animals (Fig 5A-F). These results imply that *daf-16* acts cell-non-autonomously in somatic cells in order to mediate the suppression of *prg-1*. However, we were unable to rescue *prg-1 daf-16 daf-2* sterility with *daf-16* transgenes expressed in the intestine (*ges-10* promoter, *muEx227*) or the muscle (*myo-3* promoter, *muEx212*) (Fig 4B, S1 Table) (50). Although DAF-16 has been shown to function cell-non-autonomously in the intestine or nervous system to promote somatic longevity (50), we found that expression of DAF-16 in the intestine was not capable of promoting the fertility of *prg-1* mutants. This indicates that the role of DAF-16 in repressing the fertility defects of *prg-1*/piRNA mutants is likely to be distinct from its effects on somatic longevity, which we confirmed for *daf-2; rde-4* double mutants in the accompanying manuscript (Heestand, Simon, Frenk et al).

**Fig 5.**
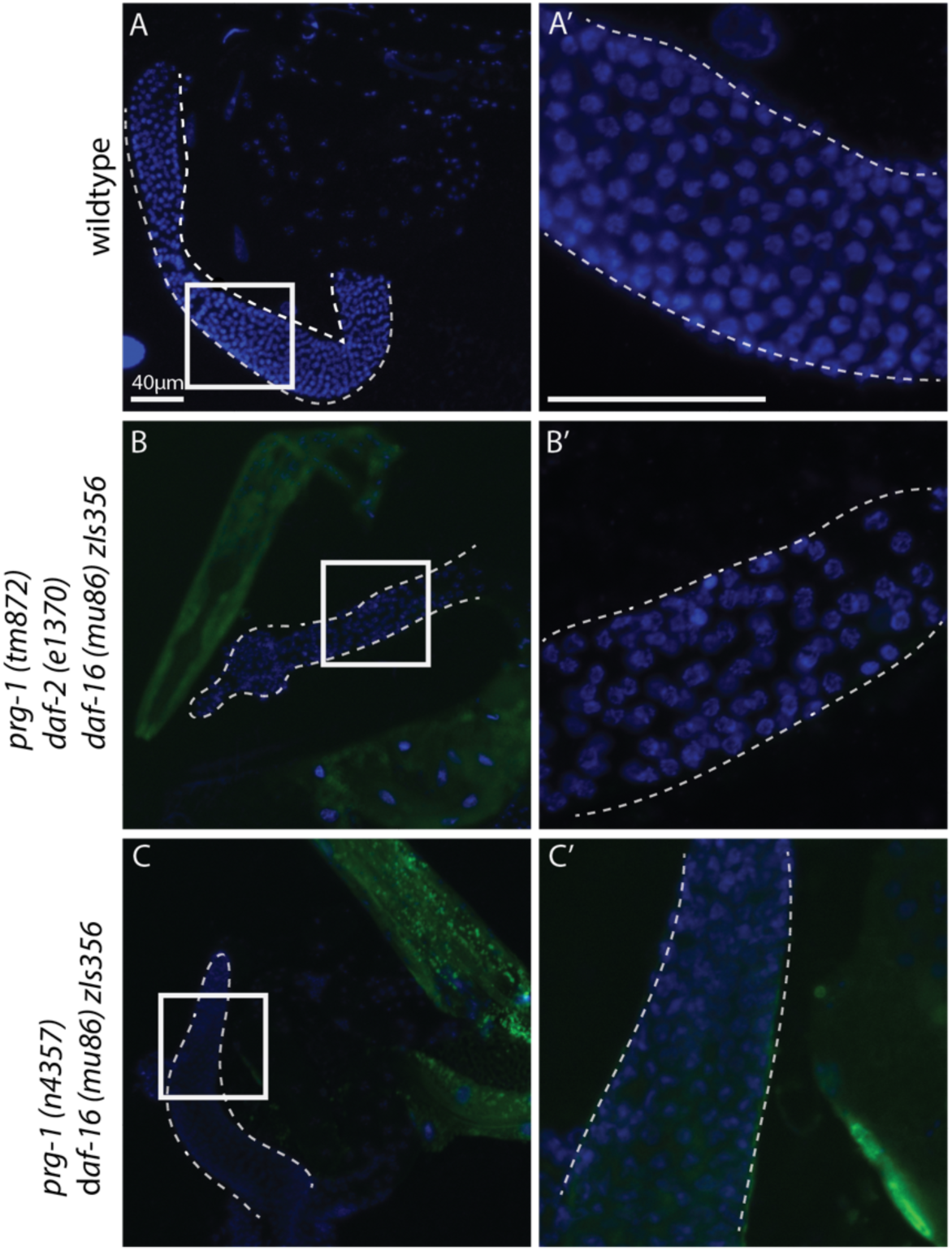
DAF-16::GFP expression in the germline. Wildtype control and two strains expressing DAF-16::GFP through the *zls356* repetitive transgene were DAPI stained and imaged after germline dissection at 10x (A-C) and 40x (A’-C’). Images show dissected germlines and parts of somatic tissue. DAPI = blue; DAF-16::GFP = green.

### DAF-16 activates a cell-non-autonomous RNAi pathway to suppress *prg-1*

We showed previously that mutation of *daf-2* eliminates the fertility defects of *prg-1* mutants by activating via the *mutator* genes *mut-7* and *rde-2*, which create 22G-RNAs that directly promote genomic silencing (12). Given that DAF-16 acts in the soma to promote fertility of *prg-1; daf-2* double mutants, we asked if DAF-16 might upregulate a systemic RNAi pathway in the soma to promote the fertility of *prg-1* mutants. The SID-1 transmembrane protein promotes systemic RNAi by importing short dsRNA fragments that are intermediates in systemic siRNA responses (29, 30, 56). We therefore constructed *prg-1; daf-2; sid-1* triple mutants, which displayed a transgenerational sterility phenotype similar to *prg-1* single mutants (Fig 4C, Table S1). These results indicate that DAF-16 acts in somatic cells to upregulate a cell-non-autonomous RNA interference pathway that suppresses the sterility of *prg-1* mutants.

The 26G-RNA pathway is an endogenous RNA interference pathway that relies on 26 nt primary siRNAs that possess a 5’ guanine (37, 38). Given the role of *sid-1* in suppression of the Mortal Germline (Mrt) phenotype of *prg-1*, we asked if genes that promote the 26G-RNA RNAi pathway were required for DAF-16 to promote fertility of *prg-1* mutants, typically by testing independent mutations per 26G-RNA pathway gene. We found that the exonuclease ERI-1 and the Argonaute protein ERGO-1, which are required for 26G-RNA biogenesis, were dispensable for suppression of *prg-1* mutant fertility defects by DAF-16 (Fig 4C, Table S1). Several proteins that promote 26G RNA biogenesis were required to rescue of the transgenerational fertility defect of *prg-1* mutants, including the *C. elegans* Dicer orthologue DCR-1, its interacting partner the dsRNA binding protein RDE-4 and the RRF-3 RNA-dependent RNA polymerase that creates primary siRNAs for the endogenous 26G RNAi pathway (Fig 6A, S1 Table) (37, 38). The exogenous RNAi pathway requires the Argonaute protein RDE-1, which promotes biogenesis of primary siRNAs from exogenous dsRNA (1), was not required to suppress the fertility defects of *prg-1* mutants (Fig 4C, S1 Table). Therefore, some but not all genes that promote the biogenesis of 26G siRNAs are required for DAF-16 to suppress the fertility defect of *prg-1* mutants.

**Fig 6.**
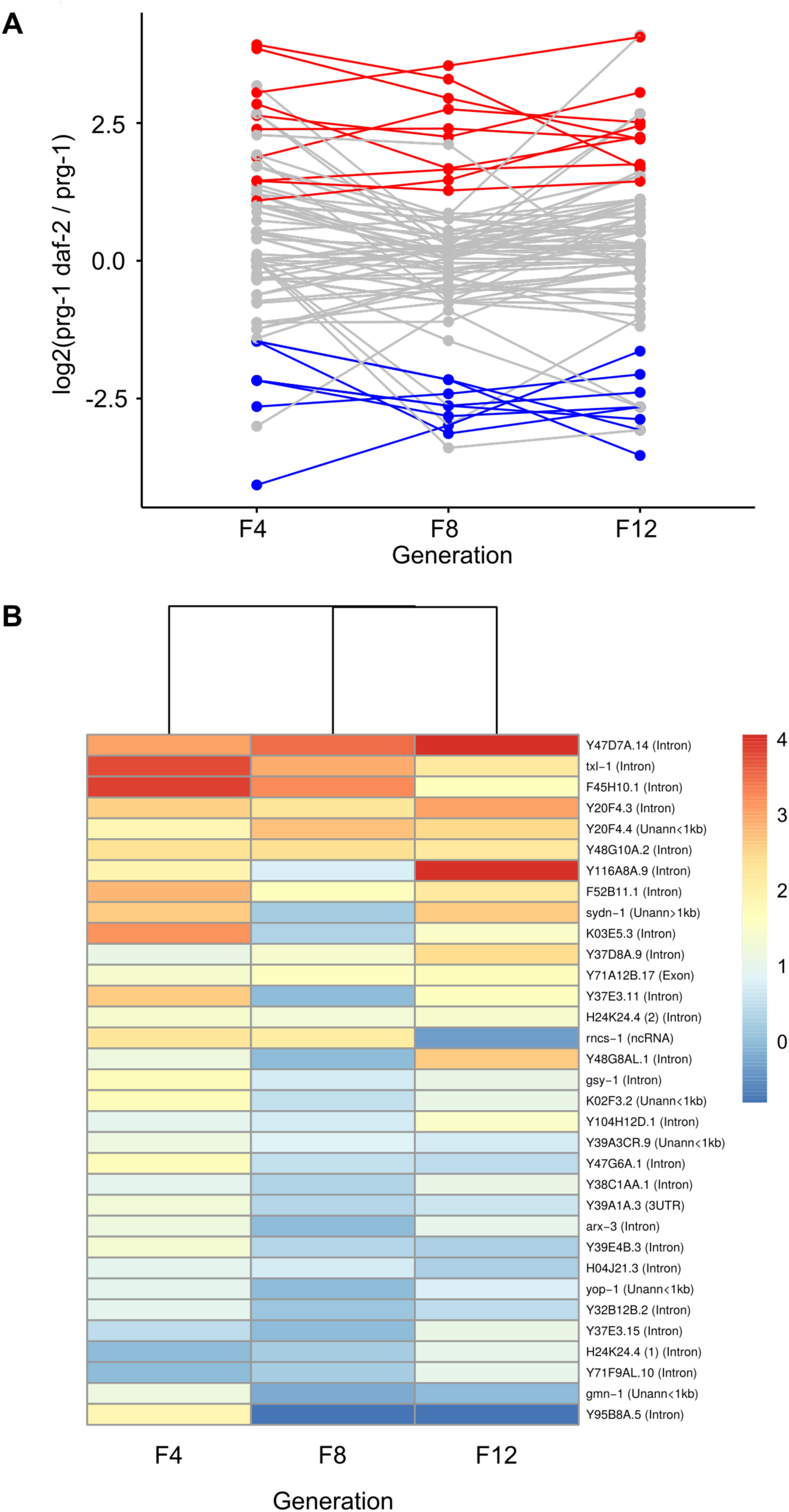
Double-stranded RNA producing loci targeted by endogenous siRNAs. (A) log2-fold change in siRNA abundance for all dsRNA loci with detectable 22Gs in *prg-1; daf-2* versus *prg-1* samples taken at different generations. Loci for which 22G RNAs are consistently upregulated and downregulated in each generation are shown in red and blue respectively. (B) Changes in relative siRNA abundance over generations for loci with increased abundance in *prg-1; daf-2* versus at least one *prg-1* sample. Color scale represents log2-fold change in *prg-1; daf-2* versus the respective *prg-1* sample.

### Analysis of dsRNA and siRNA levels in response to *daf-2* deficiency

The requirement for DCR-1 and RDE-4 in the suppression of sterility in *prg-1; daf-2* double mutants suggests the involvement of a dsRNA intermediate, because RDE-4 interacts with its targets via a dsRNA interaction domain (57). Adenosine deaminases modify double stranded RNAs by converting adenine to inosine, which results in dsRNA destruction (58) and therefore precludes dsRNA processing into 1° siRNAs by RDE-4/DCR-1 and subsequent creation of 2° 22G RNAs (59). To investigate potential *daf-2-*dependent changes in dsRNA processing, we re-analysed previously published *daf-2* RNA-seq data (60) and lists of previously defined *daf-16-* dependent loci based on RNA microarrays (61). Loss of adenosine deaminase activity results in targeting of endogenous dsRNA by the RNAi machinery (59), which is one possible mechanism by which loss of *daf-2* could drive changes in dsRNA processing. However, we found that the *C. elegans* adenosine deaminases *adr-1* and *adr-2* were unchanged in *daf-2* mutants when compared to wildtype (S3 Table), indicating that transcriptional repression of these genes is not responsible for altered dsRNA metabolism in *daf-2* mutants. We confirmed this by asking if small RNAs derived from dsRNA loci are generally upregulated by mutation of *daf-2*. We studied previously published small RNA datasets for *prg-1* mutants (generations F4, F8 and F12) in comparison with *prg-1; daf-2* double mutants (12) for changes in 22G RNA abundance at 1523 double-stranded RNA loci in the *C. elegans* genome (59). We found 33 dsRNA loci for which abundance of 22G RNAs is increased at least twofold in *prg-1 daf-2* animals compared with *prg-1* in at least one generation. 10 of these loci showed consistent 22G RNA increases for all generations (Fig. 6A, 6B). We also found 18 loci that displayed at least a twofold decrease in the levels of 22G RNAs in *prg-1 daf-2* versus *prg-1* in at least one generation, and eight loci with consistently decreased 22G RNA levels across all generations (Fig. 6A). Most dsRNAs were found to occur in introns of protein-coding genes (59). The observed differences in siRNA abundance upon loss of *daf-2* could lead to alterations in the expression of the respective genes. We looked for concordance between siRNA abundance and full-length transcript abundance by dsRNA comparing loci with differentially regulated 22Gs with a list of genes that were found to be significantly upregulated in late-generation *prg-1* but not in late-generation *prg-1 daf-2* by tiling microarray (12). Only two out of the 206 identified genes overlapped with dsRNA loci (Y69A2AL.2 and Y105C5A.13), and neither locus displayed consistently upregulated siRNAs in *prg-1 daf-2* double mutants.

Dicer has been shown to passively bind but not process some endogenous RNA targets in *C. elegans* and in human cells (62). Of the 2508 *C. elegans* DCR1-bound loci detected, 367 (∼15%) overlapped with dsRNA loci. Only three of these DCR-1-associated dsRNA loci (F45H10.1, F52B11.1 and H24K24.4) showed consistently increased siRNAs in *prg-1 daf-2* double mutants (Fig. 6B), indicating that dsRNAs that are passively bound to DCR-1 do not become actively processed in *prg-1* mutants that are deficient for *daf-2*.

*daf-2* mutants have been previously shown to be hypersensitive to exogenous RNAi, and our work with *prg-1* has defined an endogenous RNAi pathway that *daf-2* stimulates. We looked for *daf-2* dependent expression changes of genes associated with RNAi or transcriptional silencing in the *daf-2* reference data described above. Analysis of the RNA-seq data revealed a greater than two-fold downregulation of the putitative H3K79 methyltransferase *dot-1* and a greater than twofold upregulation for three histone genes (*his-12*, *his-16*, *his-43*), the putative membrane transporter *pgp-4* and *T23G4.2*, a homologue of the human mitochondrial lon pepitdase 1. However, these genes were not previously identified as DAF-16 targets (60, 61). We therefore conclude that loss of *daf-2* does not trigger strong changes to transcription of DAF-16 targets known to be involved in RNAi or transcriptional silencing pathways.

## Discussion

We found that although deficiency for *daf-2* can suppress the fertility defects of *prg-1* mutants, it was not able to suppress the transgenerational sterility defect of *nrde-1* or *nrde-4* mutants, nor of *prg-1; nrde* double mutants. This indicates that DAF-16 suppresses the sterility of *prg-1* mutants via a small RNA-mediated genome silencing response within the nucleus. This is consistent with previous results that showed that *daf-2* mutation requires secondary 22G small RNAs generated by *mut-7* and the H3K4 histone demethylase *spr-5* that promote genomic silencing (12). We studied this transcriptional silencing response to reduced insulin signalling and found that a somatic function of the NRDE pathway was required for its ability to suppress the sterility of PRG-1/Piwi. Two *C. elegans* somatic siRNA pathways have been previously described: the 26G and the anti-viral/exogenous RNAi response pathways (1). Our results indicate that neither of these pathways is responsible for suppressing the fertility defect of *prg-1* mutants. However, given that RDE-4 and DCR-1 were required, we suggest that the dsRNA-binding properties of RDE-4 mean that an endogenous form of dsRNA is the initial trigger in this DAF-16 endogenous RNAi pathway.

Based on our data, we propose the following model (Fig 7): DAF-16 activates expression of a limiting factor in an endogenous RNAi pathway that is not typically systemically activated; this could correspond to transcription or processing of a dsRNA intermediate that triggers a systemic RNAi response, or possibly to the activation or expression of a protein that promotes small RNA-mediated genome silencing in somatic cells. RRF-3 likely acts in somatic cells to create 1° siRNAs from dsRNA that associates with RDE-4/DCR-1. These primary siRNAs associate with target RNAs and the RDRP RRF-1 is recruited to generate *de novo* secondary 22G RNAs that associate with somatic NRDE-3, which may enter the nucleus to promote a nuclear RNAi response that could result in amplification of siRNAs (6) to generate siRNA-related dsRNA intermediates that are imported into germ cells by SID-1, thereby triggering a second round of 22G siRNA biogenesis that we suggest might be tertiary (30) siRNAs (Fig 7C), which have been recently observed in another context (63). However, we found that NRDE-3::GFP displayed a diffuse homogenous cytoplasmic and nuclear localization in response to mutation of *daf-2,* such that more embryos had mainly cytoplasmic or no distinct nuclear GFP::NRDE-3 expression. In contrast, loss of the dsRNA degrading adenosine deaminases *adr-1* and *adr-2* results to high levels of endogenous dsRNAs that, in *rrf-3* or *eri-1* mutant backgrounds that eliminate the endogenous 26G RNA pathway and nuclear NRDE-3 (46), are converted into siRNAs that promote strong nuclear localization of NRDE-3 (Fig. 7B) (59). Nuclear NRDE-3 can be similarly induced in response to exogenous dsRNA triggers (46). Consistent with lack of strong nuclear NRDE-3 in either *daf-2* or *prg-1; daf-2* mutants, we failed to see strong decreases in transcription of *adr-1* or *adr-2* in response to mutation of *daf-2*, nor did we observe consistent induction of siRNA levels in *prg-1; daf-2* mutants for loci known to produce dsRNA (Fig. 6). This indicates that a general alteration to dsRNA production or processing is unlikely to explain the suppression of *prg-1* sterility by *daf-2* mutation. We also found that processing of the pool of dsRNAs that are passively (constitutively) bound to Dicer (62) was not generally upregulated and we failed to identify RNAi or transcriptional silencing genes whose expression was significantly altered by mutation of *daf-2*. We suggest that one or more dsRNA loci expressed in somatic cells are likely processed and transported to the germline to become a source of siRNAs that might substitute for the lack of piRNA-mediated genome silencing in *prg-1* mutants. Interestingly, one of the most upregulated transcripts in *daf-2* mutants is the long non-coding RNA *tts-1*, which contributes to lifespan extension by binding to ribosomes and inhibiting translation (64, 65). Though siRNAs to *tts-1* were not significantly altered in *prg-1; daf-2* double mutants in comparison to *prg-1* mutants, we suggest that a similar non-coding RNA, whose transcription or processing into siRNAs may be upregulated by DAF-16, may promote germ cell immortality in *prg-1; daf-2* mutants.

**Fig 7.**
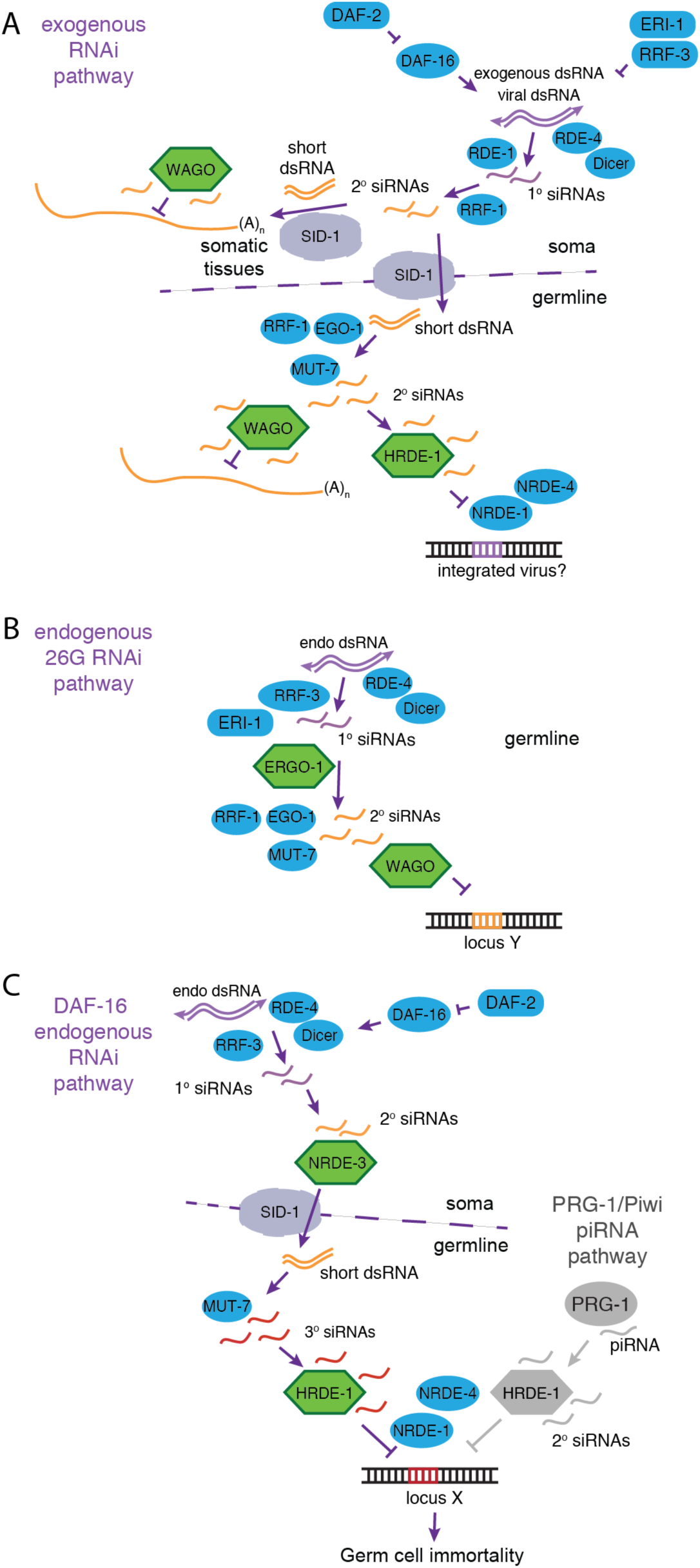
Models for exogenous and endogenous RNAi pathways. Models for (A) the exogenous RNAi pathway that can be upregulated by mutation of *daf-2* and promotes WAGO-mediated mRNA destruction or HRDE-1-mediated nuclear silencing (1, 66), (B) endogenous 26G RNAi pathway (37, 38), and (C) (this study) the endogenous RNAi pathway that is induced by mutation of *daf-2* such that somatic DAF-16 and NRDE-3 activities lead to creation of germline siRNAs (red) that interact with HRDE-1 to promote germ cell immortality of *prg-1* mutants via NRDE-1 and NRDE-4 (left side), whereas in wildtype animals PRG-1 interacts with piRNAs to promote transcriptional silencing and germ cell immortality via HRDE-1 (right side, grey color). The DAF-16-mediated endogenous RNAi pathway in panel C, left side, is distinct from the exogenous RNAi pathway in panel A because it does not require *rde-1*, and is distinct from the 26G RNAi pathway in panel B because it does not require *eri-1* or *ergo-1*.

Although we addressed several plausible mechanisms by which dsRNA metabolism might be altered in response to mutation of *daf-2*, based on known properties of the endogenous DAF-16 small RNA silencing pathway (Fig. 7C), it remains to be understood: 1) which tissue DAF-16 acts in to promote this RNAi pathway, 2) which locus or loci in somatic cells generates small RNAs that are relevant to this pathway, 3) which locus or loci in germ cells do these dsRNAs target to promote the fertility of *prg-1* mutants, and 4) if upregulation of the endogenous RNAi pathway by DAF-16 is regulated by the same mechanism that enhances the exogenous RNAi pathway in *daf-2* mutants, which protects the genome against exogenous parasites such as transposons that are transmitted by horizontal gene transfer or viruses (Fig. 7A) (66).

Systemic RNAi has been very useful as an experimental tool for suppressing gene expression, and it can suppress viral infections in plants and has the potential to suppress viral infections in *C. elegans* for multiple generations, possibly allowing small RNA-mediated transgenerational responses to both pathogenic and non-pathogenic environmental stimuli in a manner that Jean-Baptiste Lamarck and Charles Darwin envisioned might be an illustrious tool in the crucible of evolution (5, 67). Here we define a natural function of systemic RNAi (Fig 7C), where RNAi intermediates imported by SID-1 dsRNA transporter can act to suppress the transcriptional silencing defect that compromises germ cell immortality of *prg-1*/Piwi mutants. This scenario may be realistic because the PRG-1/Piwi silencing system represents an innate immune system of the germline, which might be a target of viral or transposon genomic parasites that seek to suppress the endogenous defences against their expression and replication. Viruses are well known to induce stress responses, as observed for mammalian cells that are exposed to dsRNAs (68). We posit that it is such attacks, possibly common in the arms race between host and parasite, might make the stress resistance function of the DAF-16/Foxo pathway a logical mechanism by which to activate a response to silencing of PRG-1 in a manner that ensures immortality of the germ line. In this sense, somatic diapause and germ cell immortality, which can both be promoted by high DAF-16/Foxo and low DAF-2/insulin/IGF1 receptor signalling, may represent two faces of the same coin, whose common goal is to ensure evolutionary success by strengthening the soma and the germ line in times of stress.

## Materials and Methods

### Strains

All strains were cultured at 20°C on Nematode Growth Medium (NGM) plates seeded with *E. coli* O50 (69). Strains used include Bristol N2 wild type, *prg-1(n4357) nrde-3(gg66), prg-1(tm872) nrde-3(gg66), prg-1(n4357) I, prg-1(tm872) I, nrde-1(yp4), nrde-1(yp5), nrde-4(gg131), prg-1(n4357) nrde-1(yp4), prg-1(n4357) nrde-1(yp5), prg-1(tm872) nrde-1(yp4), prg-1(tm872) nrde-1(yp5), prg-1(n4357) nrde-4(gg131), daf-2(e1368), daf-2(e1368) nrde-1(yp4), daf-2(e1368) nrde-1(yp5), daf-2(e1368) nrde-4(gg131), hrde-1(tm1200), prg-1(tm872) hrde-1(tm1200) daf-2(e1368), prg-1(tm872) nrde-3(gg66), prg-1(tm872) daf-2(m41) nrde-3 (gg66), prg-1(n4357) daf-2(e1370), prg-1(tm872) daf-2(e1370), prg-1(n4357) daf-16(mu86) daf-2(e1370), prg-1(tm872) daf-16(mu86) daf-2(e1370), prg-1(n4357) daf-16(mu86) daf-2(e1370) muls71, prg-1(n4357) daf-16(mu86) daf-2(e1370) zls356, prg-1(tm872) daf-16(mu86) daf-2(e1370) muls71, prg-1(tm872) daf-16(mu86) daf-2(e1370) zls356, prg-1(n4357) daf-16(mu86) daf-2(e1370) muEx211, prg-1(n4357) daf-16(mu86) daf-2(e1370) muEx212, prg-1(n4357) daf-16(mu86) daf-2(e1370) muEx227, prg-1(n4357) daf-2(e1368) sid-1(qt9), prg-1(tm872) daf-2(e1368) sid-1(qt9), prg-1(tm872) daf-2(e1370) eri-1(mg366), prg-1(n4357) daf-2(e1370) ergo-1(gg98), prg-1(tm872) daf-2(e1370) ergo-1(gg98), prg-1(n4357) daf-2(e1368) rde-1(ne217), prg-1(tm872) daf-2(e1368) rde-1(ne217), prg-1(n4357) daf-2(e1368) rde-4(ne301), prg-1(tm872) daf-2(e1368) rde-4(ne301), prg-1(tm872) rrf-3(pk1426) daf-2(e1368), prg-1(n4357) daf-2(e1368) dcr-1 please check, prg-1(tm872) daf-2(e1368) dcr-1(mg375), gfp::nrde-3, daf-2(e1368) gfp::nrde-3, prg-1(n4357) daf-2(e1368) gfp::nrde-3, prg-1(n4357) gfp::nrde-3*.

### Imaging and DAPI staining

DAF-16::GFP worms were dissected and fixed in 1%PFA before DAPI staining. Images were taken using immunofluorescence on Nikon Eclipse E800 microscope with a NIS Elements software. Some NRDE-3::GFP embryos were visualized as z-stacks with a LSM 710 laser scanning confocal in order to determine the total number of nuclei per embryo that was then applied to calculate the total number of embryos in single stack images. Embryos between 110-190 cell stage were selected for analysis. Images were processed using ImageJ.

### Statistical Analysis

For GFP::NRDE-3 embryo phenotypes, contingency tables were constructed and pairwise Chi Square tests were used to determine significant differences in germline phenotype distributions. For trans-generational lifespan assays, pairwise log rank tests were performed. All reported p-values were adjusted using Bonferroni correction when multiple comparisons were performed.

### Sterility Assays

Worms were grown by transferring six L1s to fresh NGM plates every week. They were defined as sterile when no more than six worms were found on the plate of the date of transfer.

### RNA-seq analysis

The following publicly available RNA-seq datasets were download from the Gene Expression Omnibus (https://www.ncbi.nlm.nih.gov/geo/): GSE40572 *(prg-1 s*mall RNAs) and GSE93724 (*daf-2* and wild-type mRNA). For mRNA-seq, adapter trimming was performed as required using the bbduk.sh script from the BBmap suite (70) and custom scripts. Reads were then mapped to the C. elegans genome (WS251) using hisat2 (71) with default settings and read counts were assigned to protein-coding genes using the featureCounts utility from the Subread package (72). For multimapping reads, each mapping locus was assigned a count of 1/n where n=number of hits. Differentially expressed genes were identified using DESeq2 and were defined as changing at least 2-fold with FDR-corrected p-value < 0.01. For small RNA-seq data, fasta files were filtered for 22G reads using a custom python script. Reads were then converted to fasta format and mapped to the *C. elegans* genome (WS251) using bowtie (73) with the following options: -M 1 -v 0 -best -strata. Reads mapping to dsRNA defined in (59) were extracted using bedtools (74). Read counts were normalized by total number of mappable reads in each library and quantified using custom R scripts. A pseudocount of 1 was added after normalization to avoid division-by-zero errors when calculating log2-fold change.

To find detect changes in genes involved in RNAi or gene silencing based on the *daf-2* RNA-seq data, a list of all GO terms was downladed from geneontology.org and filtered for GO terms matching the following regular expression: "RNAi|(RNA interference)|(siRNA)|silenc|piRNA". *C. elegans* genes matching any of the GO terms in this set were obtained from Biomart (75) and searched against the DESeq2 output from the *daf-2* mutant RNA-seq data.

## Acknowledgements

We thank members of the Ahmed lab for critical reading of the manuscript. Some strains were provided by the CGC, which is funded by NIH Office of Research Infrastructure Programs (P40 OD010440). This study was supported by NIH grant RO1 GM135470 (S.A). The funders had no role in study design, data collection and analysis, decision to publish, or preparation of the manuscript.

## Author contributions

M.S., M.S., A.H., M.G, A.N.S., A.S. performed experiments. S.F. performed RNA analysis and statistical analysis of the data. B.H., M.S., S.F. and S.A. wrote the manuscript. S.A. and A.S. supervised research.

---

**Supplemental Fig S1.**
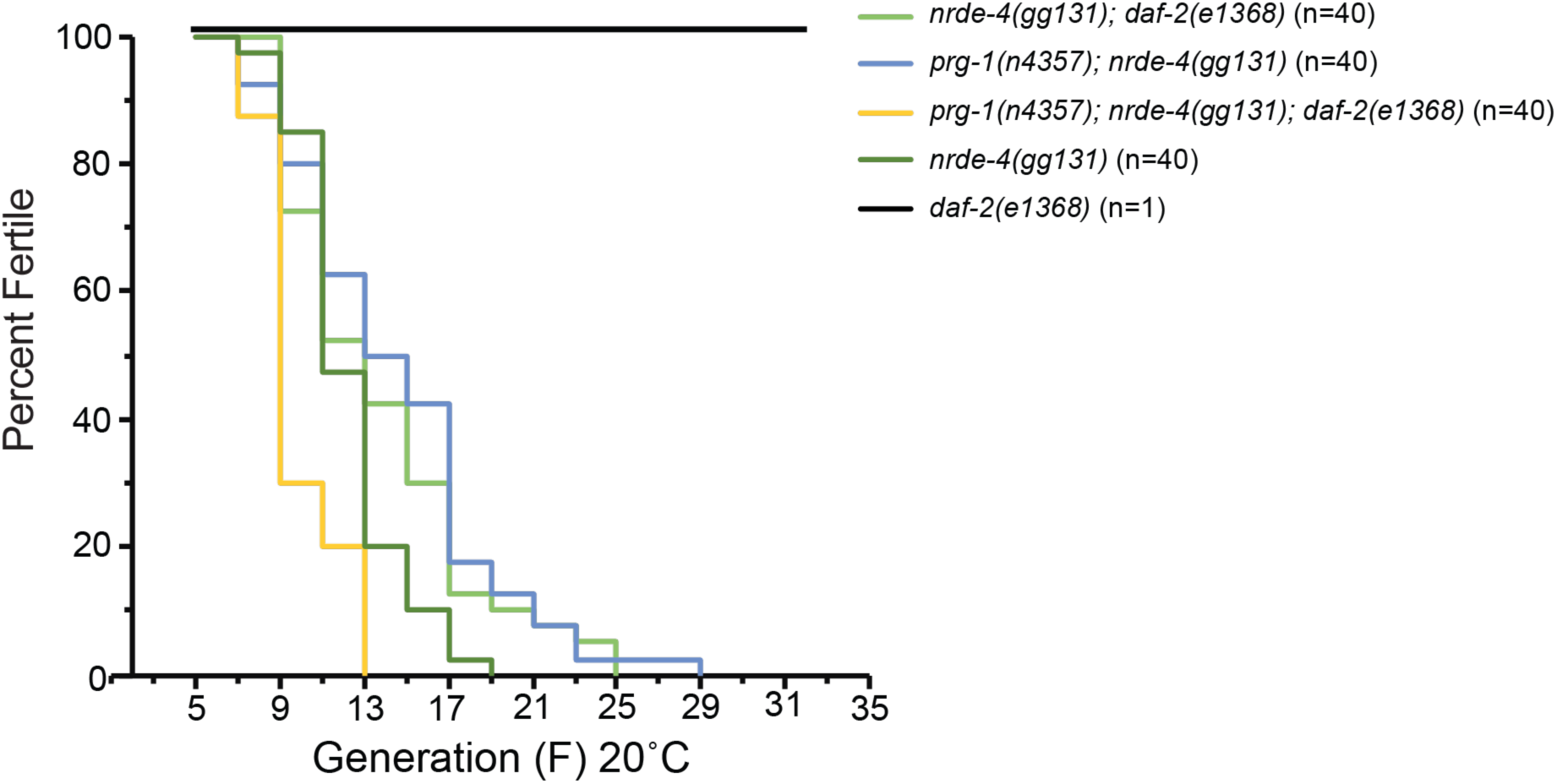
NRDE-4 is required for *daf-2* deficiency to promote fertility of *prg-1* mutants. *prg-1; daf-2; nrde-4* triple mutants display transgenerational sterility, similar to *nrde-4* and *prg-1; nrde-4* and *daf-2; nrde-4* mutants.

**Supplemental S1 Table. Germline survival for *mrt* assays.** 50% survival, maximum survival and total worms assayed for all mortal germline assays.

**Supplemental S2 Table. p values for *mrt* assays.** Log rank analysis with Bonferroni correction comparing Mortal Germline assays.

**Supplemental S3 Table. Log2-fold change and FDR-adjusted p values for genes associated with RNAi and silencing in published *daf-2* RNA-seq data.**

